# Conformational gating in ammonia lyases

**DOI:** 10.1101/583088

**Authors:** Matteo Lambrughi, Željka Sanader Maršić, Veronica Saez-Jimenez, Valeria Mapelli, Lisbeth Olsson, Elena Papaleo

## Abstract

Ammonia lyases (AL) are enzymes of industrial and biomedical interest. Knowledge of AL structure-dynamics-function relationship would be instrumental for making use of the application potential of these enzymes. We investigated, using microsecond molecular dynamics, the conformational changes in the proximity of the catalytic pocket of a 3-methylaspartate ammonia lyase (MAL) as a model system. In particular, we identified two regulatory elements in the MAL structure, i.e., the β5-α2 loop, and the helix-hairpin-loop subdomain. We showed that they undergo conformational changes switching from ‘occluded’ to ‘open’ states. We observed that these rearrangements are coupled to changes in the accessibility of the active site. The β5-α2 loop and the helix-hairpin-loop subdomain modulate the formation of tunnels from the protein surface to the substrate binding site, making the active site more accessible to the substrate when they are in an open state. We pinpointed a sequential mechanism, in which the helix-hairpin-loop subdomain needs to break a subset of intramolecular interactions first, to then allow the opening of the β5-α2 loop and, as a consequence, make the AL catalytic pocket accessible for the substrate. Our data suggest that protein dynamics need to be considered in the design of new AL variants for protein engineering and therapeutic purposes.

## Introduction

Ammonia lyases (ALs) are a broad class of enzymes which catalyze different transformations based on α- and β-amino acid scaffolds, such as the deamination and isomerization of natural amino acids through the reversible cleavage or the shifting of a C-N bond. ALs are highly heterogeneous in their structures and mechanisms of action, as attested by the fact that they cover 31 diverse EC sub-classes and they are characterized by a high stereoselectivity for their substrates [1]. These enzymes are commercially appealing for their industrial [2,3] and biomedical applications [1,2]. For example, they have been suggested as potential cancer biotherapeutics [1] due to the fact ALs could prevent the supply of the tumor cells with essential metabolites. One example is the histidine ammonia lyase which influences the growth of ovarian and prostate cancer cells by histidine deamination [1]. The deamination reaction produces urocanic acid and ammonia, eliminating histidine as building block and contributes to preventing protein synthesis, which is essential for the growth of cancer cells. The catalytic mechanism of ALs is well [3–9]. Among them, 3-methylaspartate ammonia lyases, or methylaspartases (MALs) [3] catalyze the reversible deamination of 3-methylaspartate to mesaconate. MALs belong to the enolase superfamily and they catalyze a broader range of reactions than other ammonia lyases [1] They are potentially versatile and a promising target to design new variants with different substrate specificity or improved activity and stability. For example, the MAL variant isolated from *Citrobacter amalonaticus* (CaMAL) has been engineered for enantionselective synthesis of N-substituted aspartic acids, which are essential building blocks for pharmaceutical, artificial sweeteners, synthetic enzymes and peptidomimetics [10,11].

MAL is a dimeric enzyme. Only a few crystallographic three-dimensional (3D) structures of the dimeric form of MALs are available, deposited in the Protein Data Bank (PDB), as the entries 1KKO, 1KKR [12], 1KD0, 1KCZ [13], along with two mutated variants with PDB entries 3ZVH, 3ZVI [11]. CaMAL is a homodimeric enzyme where each monomer (here referred to as A and B) is composed by 826 amino acids and can be divided into two domains: an N-terminal (residues 1–160) and a C-terminal domain (residues 170–413) connected by β-sheet regions (**Figures 1A-B**). The C-terminal domain folds into a triosephosphate-isomerase (TIM) barrel structure, consisting of eight α-helices and eight parallel β-strands, while the N-terminal domain is composed of a three antiparallel β-strands and four α-helices (**Figure 1B**).

**Figure 1.**
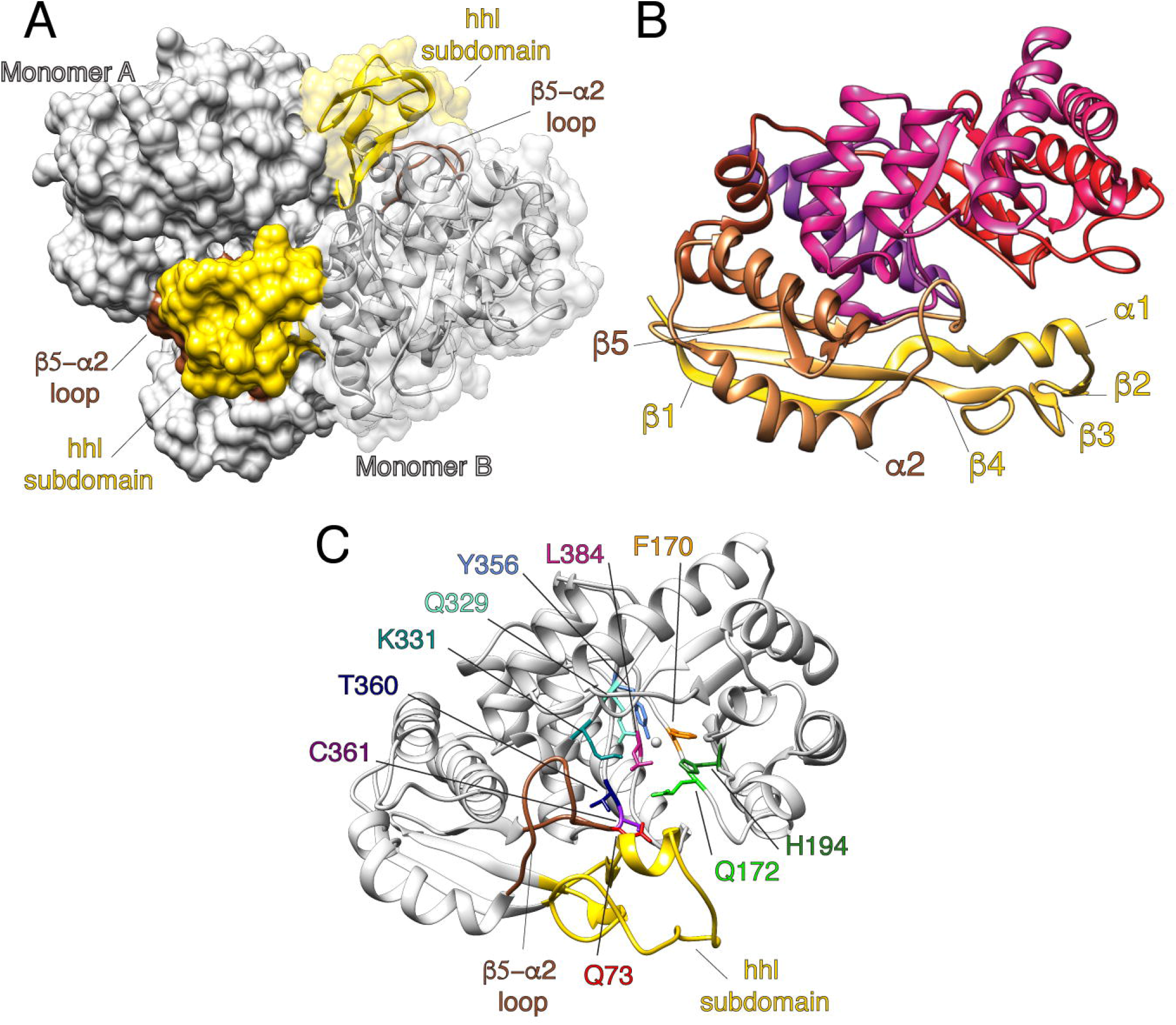
The 3-methylaspartase ammonia lyase from *Citrobacter amalonaticus* (CaMAL) has two structural elements in the proximity of the catalytic pocket, i.e., the helix-hairpin-loop (hhl) and the β5-α2 loop. A) The homodimeric structure of CaMAL (PDB entry 1KKR) is shown as white surface and the monomer B is shown as cartoon for sake of clarity. The hhl subdomain and the the β5-α2 loop are in gold and brown, respectively. B) The monomer A of CaMAL dimer is shown as cartoon and represented with a gradient of color from N-terminal (brown) to C-terminal (purple) with indicated the secondary structural elements in the proximity of the hhl subdomain (α1, β1-β4) and the β5-α2 loop (α2 and β5). C) The residues in the catalytic pocket are shown as sticks with the following color coding: Q73 red, F170 orange, Q172 green, H194 forest green, Q329 aquamarine, K331 dark cyan, Y356 cornflower blue, T360 navy blue, C361 purple and L384 violet red. The hhl subdomain and the β5-α2 loop are colored in gold and brown, respectively.

The catalytic mechanism of CaMAL has been inferred from the analysis of its 3D structure, the comparison with other members of the enolase family and validated by experimental mutagenesis [14,15]. The mutagenesis studies revealed the importance of specific residues on the structural integrity, activity, and regio- and diastereoselectivity of CaMAL (**Figure 1C**). The reaction catalyzed by CaMAL requires the presence of cations, generally provided by the metal magnesium (Mg^2+^) atom in the catalytic pocket, while the residues K331, H194 and Q329 form the catalytic triad (**Figure 1C**). K331 acts as the base catalyst, while the Mg^2+^ metal ion and the residues H194 and Q329 are responsible for the stabilization of the enolate anion, and binding to the 4-carboxylate group of the substrate [14]. Additionally, Q73, F170, Q172, Y356, T360, C361 and L384 interact with different functional groups of the substrate and are located in the pocket (**Figure 1C**).

The N-terminal domain of CaMAL bears, in the proximity of the catalytic pocket, two structural elements, that we refer as β5-α2 loop (residues 70-85) and helix-hairpin-loop subdomain (hhl, residues 12-51) (**Figure 1C**). The β5-α2 loop connects the strand β5 with the helix α2 and includes Q73, which form water-mediated hydrogen bonds with the substrate and alters CaMAL k_cat_ when mutated [15]. The hhl subdomain contains a short α-helix (residues 19-25), a small β-hairpin (β2-β3 strands, residues 29-34) and the loop between the strands β3 and β4. The hhl subdomain is involved in intermolecular interactions at the interface between the monomers [12].

At the best of our knowledge, there is no information available on the conformational changes of MALs that could be related to its function. It is a common feature of several enzymes that disordered or partially structured regions of a folded protein undergo conformational changes that modulate the access of the substrate to the active site or allow for its formation [8,16,25,17–24]. A detailed investigation of the dynamics of MALs is an essential step to understand how to engineer it for applicative purposes.

We focused on CaMAL, presenting the first all-atom Molecular Dynamics (MD) investigation of this family of enzyme and its implications for activity. Biomolecular simulations are useful to describe protein functional dynamics, from local to more global motions and the notions of the importance of dynamics for enzyme function is well established [26–31]. This includes local changes in the proximity of the active or substrate binding sites [25,32–37] up to conformational changes of large amplitude associated with the opening and closing of gating loops or domains that can modulate the access to the catalytic site or the substrate binding pocket of an enzyme [16,22,38].

## Results

### CaMAL is a dimer in solution

To select which form of CaMAL to study with MD simulations, we investigated the preferred quaternary structures of CaMAL in solution. In particular, we performed Size-Exclusion Chromatography (SEC) to estimate CaMAL molecular weight (MW). We used proteins with a known MW as calibration standards (see Materials and Methods). The resulting partition coefficients (K_av_) plotted against the logarithm of MW were fitted with a linear function (correlation coefficient ≈ 0.99) (**Figure 2A**). We identified one single sharp peak for CaMAL, corresponding to a MW of ≈ 80.4 kDa (**Figure 2A**), which is approximately double of the MW estimated for the monomeric form of CaMAL (from its primary sequence, i.e., 45.5 kDa). We concluded that CaMAL is mostly a homodimer when in solution in its substrate-unbound state, in agreement with what previously reported [39].

**Figure 2.**
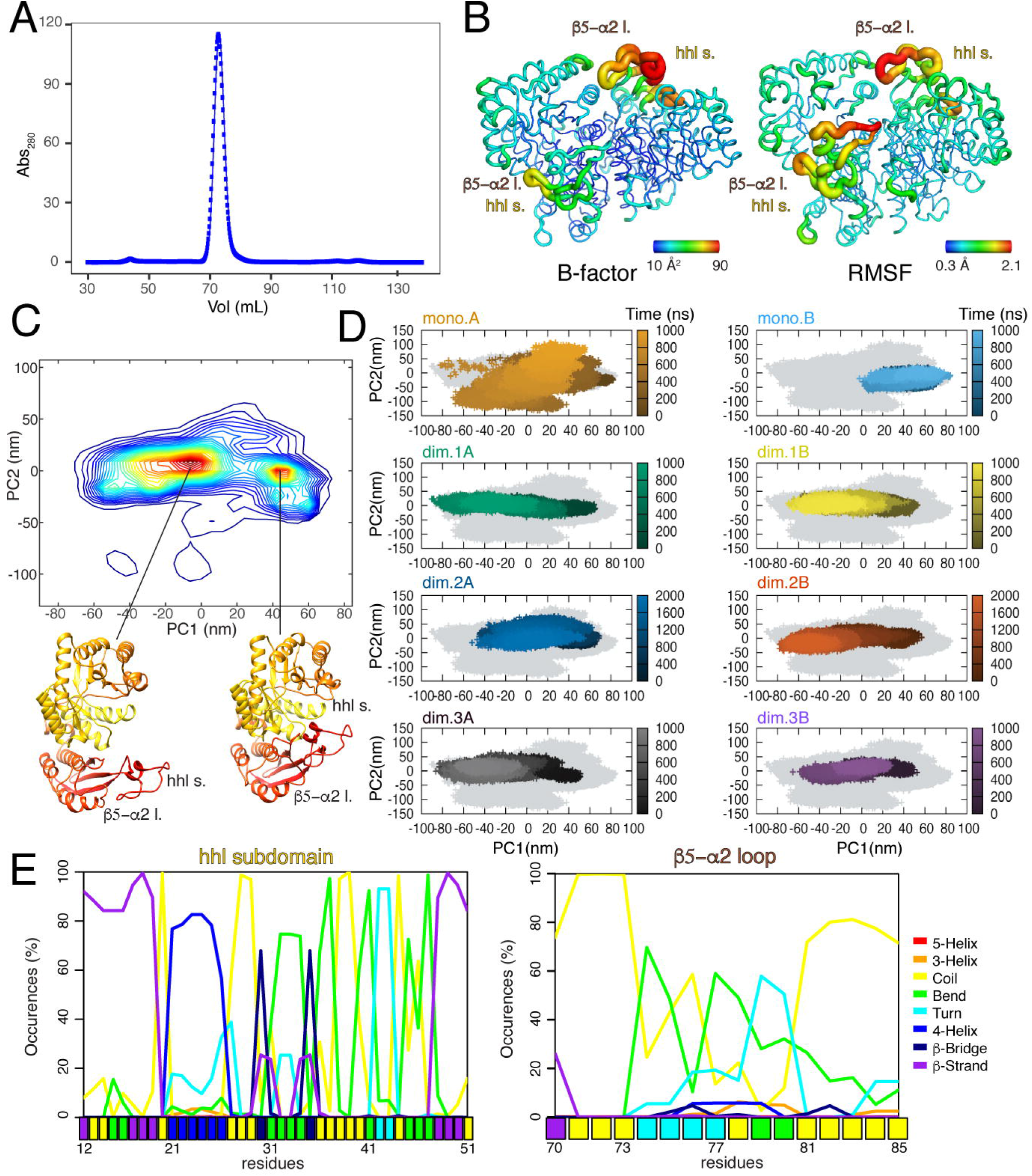
CaMAL is a dimer in solution with two highly flexible regions. A) Size exclusion chromatography experiment shows single absorption peak at the molecular mass of 80.4kDa which corresponds to the dimeric form of CaMAL. B) The B-factor (in the left) and RMSF (in the right) values are shown as tube representation on the structure of CaMAL dimer. The thickness and the color spectrum of the tube are proportional to the flexibility of the residues, with thicker red tubes representing highly flexible regions and thinner blue tubes indicating more rigid regions of the enzyme. The highest B-factor and RMSF values can be observed in the hhl subdomain and the β5-α2 loop. C) 2D distribution plot calculated using as reaction coordinates the first and the second principal components from the PCA analysis performed on all the protein atoms. We identified two densely populated subsets of conformations (indicated as ‘open’ and ‘closed’ states). One representative structure from each state is shown as cartoon in the lower part of the panel for illustrative purposes. The ‘closed’ structural ensemble included conformations of the CaMAL monomers similar to the initial X-ray structure. The ‘open’ ensemble accounted for conformations of the CaMAL monomers with both the β5-α2 loop and the hhl subdomain in open states. D) We highlighted the PCA subspace sampled by each trajectory. For the MD simulations of the dimer we showed the monomer A and B separately for sake of clarity. Simulations of the monomers and dimers are indicted as ‘mono’ and ‘dim’, respectively and the MD replicates of the dimeric form are numbered sequentially from one to three. The time evolution of each trajectory in the conformational space is represented as shade of color, from dark (0 ns) to light (end of the trajectory). We observed the transition from open to closed states in the dimer simulations. The differences observed for the simulations of the monomer forms are discussed in the text. E) We estimated the occurrence of different secondary structural classes in the regions of the hhl subdomain and β5-α2 loop during the simulations. The color bars indicate the structural classes in the X-ray structure, as a reference. For sake of clarity only data obtained for the first replicate of CaMAL dimer are reported. The other results are in the Github repository associated to the publication.

In light of the SEC results, we focused on the CaMAL dimer in solution for our MD simulations. We also collected two additional simulations of the monomers (i.e., chains A and B from the dimer structure) as a control.

### CaMAL shows asymmetric dynamic patterns with high flexibility located at the β5-α2 loop and the hhl subdomain

We employed all-atom explicit solvent one-μs MD simulations to investigate CaMAL dynamics-structure-function relationship. In particular, we aimed at identifying conformational changes that modulate the accessibility of the active site. We calculated the average per-residue Root Mean Square Fluctuation (RMSF) of the Cα atoms for each monomer of the CaMAL dimer as a measure of protein flexibility. The peaks in the RMSF profile indicate the protein regions interested by the highest flexibility (**Figure 2B**). We observed marked fluctuations in the β5-α2 loop and the hhl subdomains in both the monomeric and dimeric states of CaMAL. We found similar fluctuations also in a MD simulation performed starting from another crystallographic structure of CaMAL dimer (PDB entry 1KK0), supporting our findings (see Github repository for more details).

The simulations performed on each separated monomer show higher flexibility in the hhl subdomain in the case of the monomer A than in the dimer, while in the monomer B this region is more rigid (**Figure S1**). These differences suggest an asymmetry in the structure and dynamics of the two monomeric subunits of CaMAL, a trait common to other dimeric enzymes [40–43] (see GitHub repository for more details). This asymmetry is observed for both the hhl subdomain and the β5-α2 loop, indicating that the dynamic patterns of these regions, in the dimer, are modulated by the presence of the partner monomer. The RMSF estimated by our MD simulations is also in agreement with the crystallographic B-factor values associated with the experimental structures of the CaMAL dimer in its substrate-bound (PDB entry 1KKR) or -unbound (PDB entry 1KKO) states (**Figure 2B**).

### The β5-α2 loop and the hhl subdomain populates ‘open’ and ‘closed’ states in the μs timescale

Dimensionality reduction techniques help to identify the most important conformational transitions in a MD ensemble and compare simulations [44–47]. In particular, we employed Principal Component Analysis (PCA) [48] and focused on the ‘essential subspace’ described by the first two principal components (PCs) (**Figure 2C-D and Figure S2**). We identified two main conformational states, associated with conformational transitions of the β5-α2 loop and the hhl subdomain (**Figure 2C**). The two regions can populate ‘open’ or ‘occluded/closed’ conformations with respect to the active site (**Figure 2C**) with a progressive opening during the simulation time (**Figure 2D**, color gradient).

We observed a different dynamic pattern for the isolated monomer A with a large exploration of the PCA subspace during the simulation time (**Figure 2D**, orange gradient), which is related to its enhanced flexibility (**Figure S1**). On the contrary, CaMAL remains in its closed state in the MD simulations of the isolated monomer B, mostly sampling conformations around the starting structure (**Figure 2D**, light blue gradient), in agreement with the lower RMSF observed for this trajectory (**Figure S1**).

Moreover, we extended the second replicate of the CaMAL dimer to two μs since in this replicate the opening of the catalytic pocket was occurring after hundreds of ns (**Figure 2D**, blue and dark orange gradients).

We then analyzed more in detail the structural changes, which were associated with the conformational transitions of in the β5-α2 loop and the hhl subdomain (**Figure 2E**). The opening of the β5-α2 loop were associated with instability of the β-hairpin (residues 29-34), loss of β structure in the β2 and β3 strands (approximately in 75% of the simulation frames) and changes to bend/coil conformations (**Figure 2E**). The hhl subdomain was stable during the conformational transition, with the residues 19-25 in a stable helical conformation in all the simulations (approximately in more t1 han 70% of the simulation frames) (**Figure 2E**).

### The hhl subdomain and the β5-α2 loop modulate the accessibility of the catalytic pocket of CaMAL with a sequential mechanism

The conformational transitions observed during CaMAL simulations involved significant rearrangements of the β5-α2 loop and hhl subdomain in comparison to their position in the starting structure (**Figure 3A**). To understand the relationship between these rearrangements and changes in other regions of CaMAL, we monitored, over the simulation time, representative pairwise distances between Cα atoms of each of the residues of the β5-α2 loop or of the hhl subdomain and the Cα Q329 residue, used as a reference of the pocket (**Figure 3B**). We then estimated the pairwise Pearson correlations of the selected distances with the solvent accessibility of the catalytic pocket of CaMAL, designing a ‘correlogram’ where only correlations with a p-value lower than 0.01 are highlighted (**Figure 3C**). We calculated the solvent accessible surface area (SASA) of each of the residues of the pocket (i.e., Q73, F170, Q172, H194, Q329, K331, Y356, T360, C361 and L384), along with their cumulative SASA and the total protein SASA for each monomer. The correlogram turned out to be an efficient approach to investigate possible couplings between changes in the residues of the β5-α2 loop or the hhl subdomain and the accessibility of the catalytic pocket of the enzyme. We observed statistically significant positive correlations between the total SASA of the pocket and the distances between Q329 and many of the residues of the β5-α2 loop and the hhl subdomain (**Figure 3C**). Moreover, we applied a hierarchical cluster approach on the correlation matrices and we identified two main groups of structural measurements with higher positive correlations between them (**Figure 3C**). The first group includes the distances with the residues of the β5-α2 loop and the residues 12-14, 16, 29-33 and 48-51 of the hhl subdomains that are correlated with both the total SASA of the pocket and the accessibility of K331 and T360. These results suggest that the opening of the β5-α2 loop modulates the accessibility of the binding pocket directly, acting of the residues that are located at its entrance, such as K331 and T360 (**Figure 3B**). The second group includes the distances with the residues 17-28 and 34-47 of the hhl subdomains that are correlated with the total protein SASA and the accessibility of Q73. Q73 is located in the β5-α2 loop, and our analysis suggests that the hhl subdomains play a role in acting as a conformational gate for the β5-α2 loop (see below **Figure 4D**).

**Figure 3.**
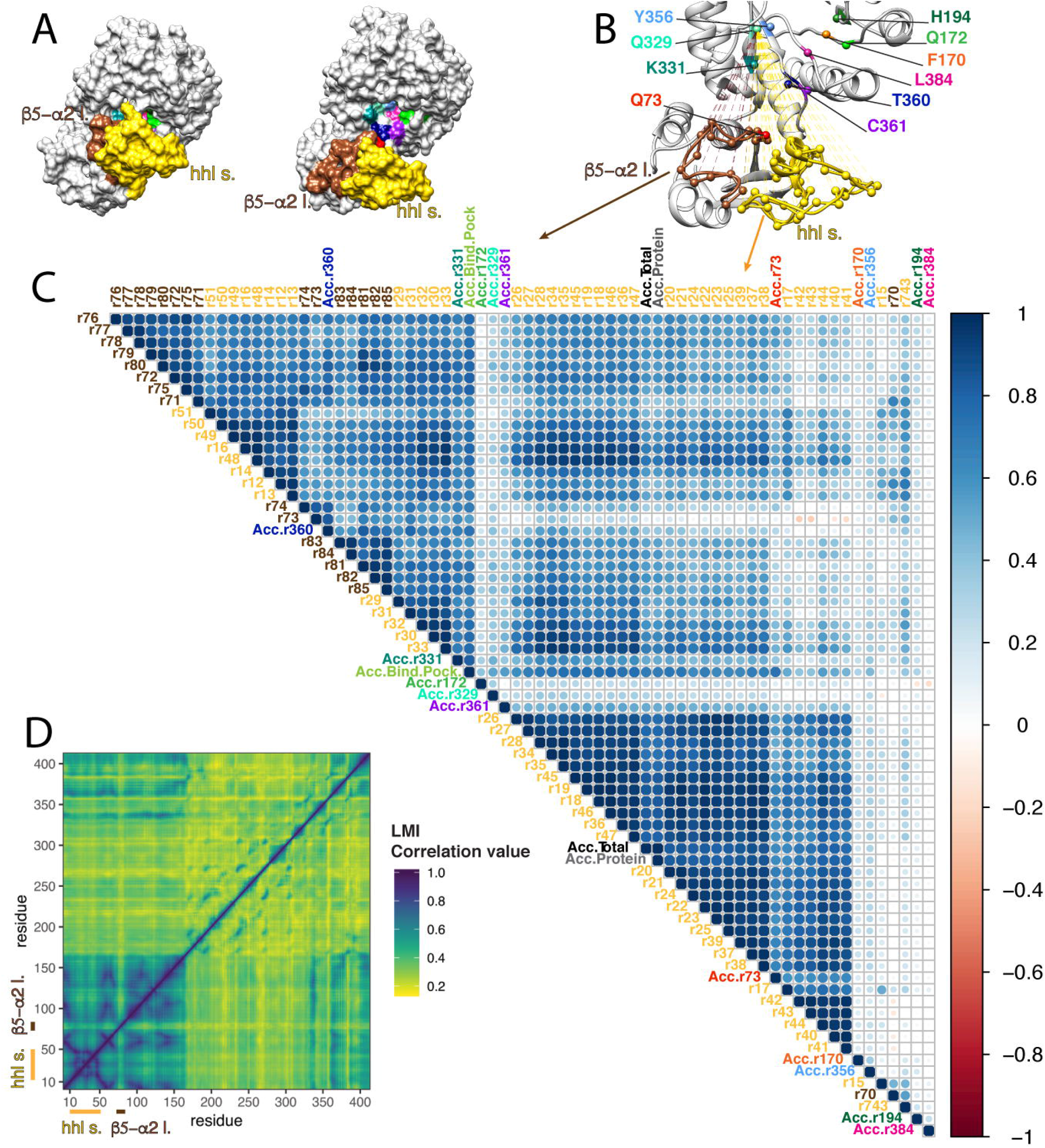
The opening of the hhl subdomain and the β5-α2 loop is correlated with changes in the accessibility of the catalytic pocket of CaMAL. The residues of the catalytic pocket are shown with the following color coding: Q73 red, F170 orange, Q172 green, H194 forest green, Q329 aquamarine, K331 dark cyan, Y356 cornflower blue, T360 navy blue, C361 purple and L384 violet red. The hhl subdomain and the β5-α2 loop are showed in gold and brown, respectively. A) Surface representation of two representative structures belonging to the ‘closed’ (in the left) and ‘open’ (in the right) states of the enzyme. The open conformations promote a higher accessibility of the residues in the catalytic pocket. B) We showed the distances between the Cα atom of Q329 in the pocket and the Cα atoms of all the residues in the β5-α2 loop and in the hhl subdomain that we used in the analyses. The CaMAL is shown as white cartoon and the residues in the binding pocket, for which the solvent accessibility surface area (SASA) was calculated, are shown as sticks. C) The correlogram shows the Pearson correlation coefficients between each pair of measurements (i.e., the selected distances and SASAs) at a p-value < 0.01. The color intensity and the size of the circle are proportional to the correlation coefficients, i.e. the strength of the correlations, between each pair of measurements, from dark red (negative correlations) to dark blue (positive correlations). The correlogram shows a clear coupling between changes in the the β5-α2 loop and in the hhl subdomain and accessibility of specific residues of the catalytic binding pocket. The details are discussed in the text. D) The heatmap shows the average correlated motions in the MD trajectories, using Linear Mutual Information (LMI). The LMI correlations are shown as a gradient from 0 (i.e., uncorrelated motions shown in yellow) to 1 (i.e., fully correlated motions in dark blue). The LMI correlation matrix shows a coupling in the motions of the β5-α2 loop and the hhl subdomain, which are also coupled to the entire N-terminal domain. For sake of clarity, we reported the results for the monomer A of the first MD replicate of the CaMAL dimeric form. The other results are in the Github repository associated to the publication.

**Figure 4.**
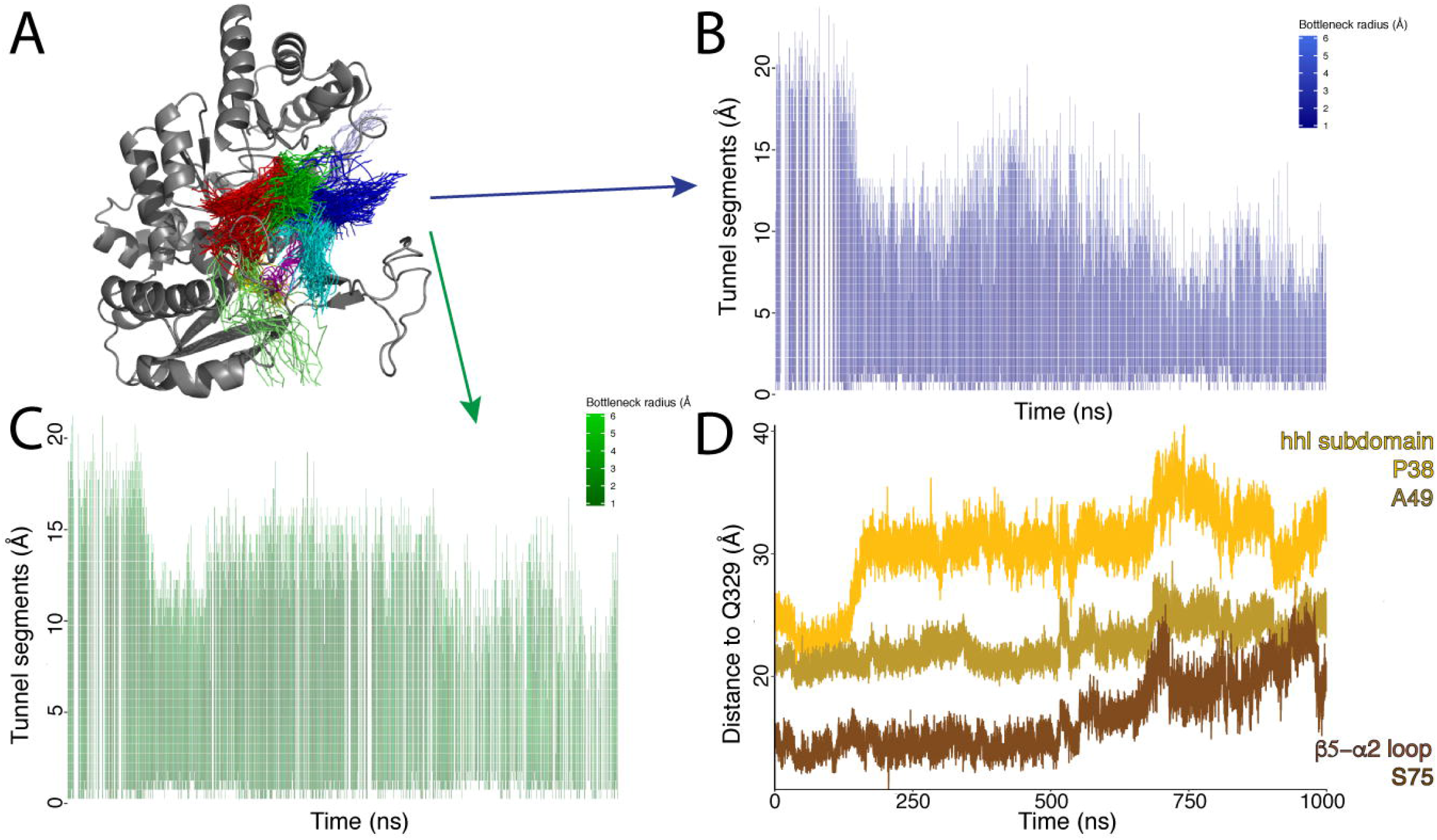
The opening of the β5-α2 loop and hhl subdomain modulate the length and the apperance of tunnels toward the active site of CaMAL. A) We showed the main clusters of tunnel pathways calculated from the MD ensembles on the structure of the CaMAL monomer. In particular, the tunnels calculated for each conformation are shown as pathway centerlines and grouped into clusters. For each conformation, if more than one tunnel pathway is grouped in the same cluster, only the pathway with the lowest cost is shown. B-C) We showed, as heatmaps, the temporal evolution of the two main cluster of tunnels in blue and green, respectively. The plots report the average length and the radius (colors in the plot) of the tunnels in the cluster for each trajectory frame. For sake of clarity, we illustrated only the results for two clusters for the monomer A of the first MD replicate of the CaMAL dimer form. The other results are in the Github repository associated to the publication. D) We showed the temporal evolution of three distances between the Cα atoms of Q329 (in the pocket) and S75 (in the β5-α2 loop, in brown), P38 and A49 (in the hhl subdomain, in gold and dark gold, respectively). The origin of the y-axis is set to the lowest value observed for the three selected distances (10.77 Å). We showed only the results for the monomer A of the first MD replicate of the CaMAL dimer form for sake of clarity. The other results are in the Github repository associated to the publication.

In addition, we observed coupling between several residues of the β5-α2 loop and the hhl subdomain (**Figure 3C**). Since these two structural elements are localized spatially close to each other in the CaMAL structure and they form intramolecular contacts, we estimated if their motions are significantly coupled at the level of residue-residue displacements. In particular, we calculated the residue-residue correlated motions along the trajectories using Linear Mutual Information (LMI, [49]). We calculated the average LMI correlation matrices over time windows of 10 and 100 ns for the MD simulations of CaMAL dimer to assess two different timescales (**Figure 3D**). We calculated the pairwise differences between the LMI matrices calculated for each of the two monomers in the same replicate, along with the differences between the LMI matrices for the monomers A of different replicates (**Figure 3D** and **Figure S3**). We concluded that the LMI matrices described similar patterns of correlated fluctuations in the monomers, despite their asymmetry (see above) with differences in the range of +/−0.3. We also observed modest differences among LMI matrices averaged over different time windows. Once assessed the robustness of the LMI matrices, we analyze them to identify local couplings in the MD ensemble (**Figure 3D**). In particular, we observed correlations in the motions of the β5-α2 loop and hhl subdomain (**Figure 3D**). Moreover, we identified coupled motions between these two structural elements and most of the N-terminal domain of CaMAL (i.e., residues 1-163), suggesting that the entire N-terminal domain undergoes conformational changes relatively to the C-terminal domain.

In summary, we observed couplings between the motions of the β5-α2 loop and the hhl subdomain, along with couplings of these two regions with the accessibility of the catalytic site. We also observed that the opening of the hhl subdomain occur before the opening of the β5-α2 loop in each replicate. Our results thus suggest that the hhl subdomain play a role in constraining the motions of the β5-α2 loop, acting as a conformational gate.

### The opening of the β5-α2 loop and hhl subdomain shorten the tunnels from the protein surface to the active site of CaMAL

We estimate the formation and time-evolution of tunnels towards the catalytic site (**Figure 4**). We used the position of four residues of the active site (i.e., Q329, D238, L384, Y356) as a reference point for the tunnel analysis. We calculated the evolution of the solvent-accessible tunnels from the protein surface to these residues with a probe radius of 1.5 Å, which resembled the size of a water molecule. The analysis identified from eight to ten clusters of tunnels depending on the trajectory (**Figure 4A**). We calculated the time evolution of the average length and radius of the tunnels in each cluster, which indicate the absence of bottlenecks when a tunnel was identified (**Figure 4B-C**). We observed that the length of the tunnels decreases with the opening of the hhl subdomain and the β5-α2 loop, showing that binding pocket became more easily accessible (**Figure 4B-C**). We also performed the same analysis with a probe radius of 2.0 Å to better resemble the size of the natural substrate (i.e., 3-methylaspartate), obtaining similar results (see Github repository). In summary, our results indicate that the opening of the hhl subdomain occur first (**Figure 4D**) and allow the β5-α2 loop outward displacement. Moreover, their concerted opening provides an efficient access to the active site, shortening the protein tunnels from the protein surface to the catalytic residues.

### Complex networks of contacts modulate the conformational changes from ‘closed’ to ‘open’ states of CaMAL

To dissect the mechanism of opening of the substrate binding pocket through the conformational changes of the hhl subdomain and the β5-α2 loop, we also used an approach based on contact maps [50]. We measured pairwise intramolecular contacts between residues, their persistency along the trajectory time and estimated the evens of breaking and formation of the interactions. We observed that the residues of the β5-α2 loop and the hhl subdomain in the open states are generally involved in a lower number of contacts at a lower persistence then in the closed states (see Github repository).

A cluster analysis based on the RMSD between the contact maps along the trajectory identified five sequential clusters (**Figure 5A**). The clusters allowed to study the temporal evolution of the contacts along the trajectories during the transition from closed to open states of the β5-α2 loop and the hhl subdomain. We estimated the percentage increase in the pairwise distances between the residues in the β5-α2 loop and the hhl subdomain with the ones in the rest of the protein, comparing the average contact maps of the first and fifth cluster. We identified several pairs of residues for which we observed a marked increase in the distances along the simulation time. For these residues, we measured the distances with respect to residues of the β5-α2 loop (**Figure 5B**) and the hhl subdomain (**Figure 5C**), as distance networks.

**Figure 5.**
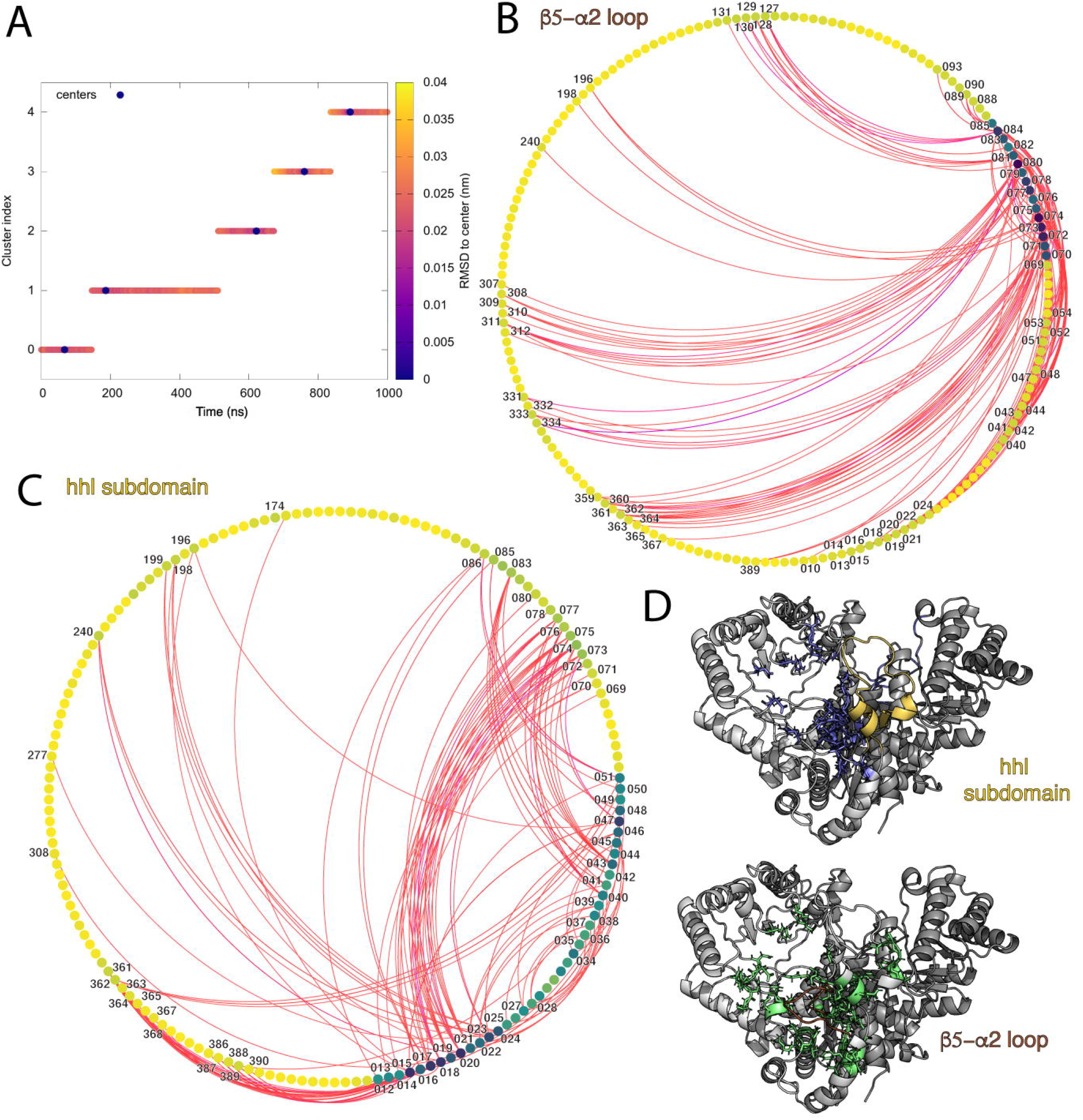
Rearrangements of intramolecular contacts during the opening of the hhl subdomains and the β5-α2 loop. A) We performed a cluster analysis based on the RMSD between the contact maps collected at different time points during the simulation, setting the maximum number of clusters as five. The plot shows the time evolution of the identified five clusters with dots colored according to the distance (measured in RMSD) of each conformation from the cluster medoid, which is indicated with a blue dot. B-C) Circular plots showing the pairwise distance network for the β5-α2 loop (B) and the hhl subdomains (C). We estimated the percentage increase of the pairwise distances between the residues of the β5-α2 loop and the hhl subdomain with the residues of the rest of the protein, comparing the average contact maps of the first and fifth cluster. In the graphs, the edges indicate percentage increase above 50% in the network and connect pairs of residues, which represent the nodes. The values of percentage increase are colored as a gradient from red (50%) to purple (300%). The node corresponding to each residue are shown as points and they are colored on the base of their degree in the network, from low (yellow) to high (dark blue). D) We mapped the network of residues on a representative open conformation of the CaMAL dimer as green and blue sticks for the β5-α2 loop and the hhl subdomain, respectively. The hhl subdomain and the β5-α2 loop are highlighted in gold and brown, respectively and both the monomers are shown. For sake of clarity, we illustrated the results for the monomer A of the first replicate of the CaMAL dimeric form. The other results are in the Github repository associated to the publication.

The distance network analysis for the β5-α2 loop (**Figure 5B**) showed a significant increase of distances with residues at both side of the entrance of the pocket: L196, N198, Y240, M389, G359-S367 (containing two catalytic residues, i.e., T360 and C361) on one side and, D307-T312 and K331-D334 (containing the catalytic residue K331), on the other side. Moreover, we observed significant changes in the distances with amino acids at the bottom of the loop, such as H88-I90, L93, and H127-R131, along with several residues in the hhl subdomain or its proximity, as S13-Y16, D18-A22, K24, T40-T44 and E51-S54 (**Figure 5B**).

The distance network for the hhl subdomain (**Figure 5C**) showed an increase in the distances with residues around the entrance of the pocket, such as for G174, L196, N198, N199, Y240, D277, E308, C361-E365 (containing the catalytic residue C361), S367, A368, and K386-G390. Furthermore, significant increases in the distance network are associated with the β5-α2 loop, such as the residues D69-G78, R80, L83, L85, and A86.

The mapping of the residues interested by changes in the distance network during the opening of the hhl subdomain and the β5-α2 loop (**Figure 5D**) showed that the hhl subdomain is involved in modulating a large area of the protein, while the β5-α2 loop is mostly associated with the regions around the active site. In summary, the contact-based distance network analysis allowed to identify the most significant molecular changes associated with the opening of the hhl subdomain and the β5-α2 loop.

## Discussion

An essential step toward the application of enzymes to industrial processes, protein engineering and biomedical treatments, is to understand their structural properties, dynamics and unveil the link with enzyme function and activity. ALs have been proposed as therapeutic agents against cancer, but despite their potential, low stability and rapid clearance hindered their applications [1]. The development of new modified variants of ALs will open up for broad applications, but it requires more detailed knowledge on their structural properties and functional mechanisms. We contributed to fill this gap using computational structural biology. Our study highlighted two important regulatory elements for CaMAL, i.e., the β5-α2 loop and hhl subdomain. They are highly flexible and undergo conformational changes on the μs timescale, switching from closed to open states with respect to the catalytic site. The hhl subdomain and the β5-α2 loop modulate the accessibility of the catalytic pocket of the enzyme, shortening the tunnels to access the active site, in a sequential mechanism where hhl subdomain acts as a conformational gate and needs to open first to promote the detachment of the β5-α2 loop from the residues at the entrance of the pocket. This mechanism might have broader relevance for ALs, as attested by the fact that other similar modulators have been reported, for example in aspartate, phenylalanine and histidine ammonia lyases, [4,8,18] These ALs do not share similarities in sequence and are structurally unrelated but, the presence of loops acting as opening/closing ‘clamp’ above the active site pinpoint interesting similarities with the molecular mechanisms that we identified in CaMAL (**Figure 6**).

**Figure 6.**
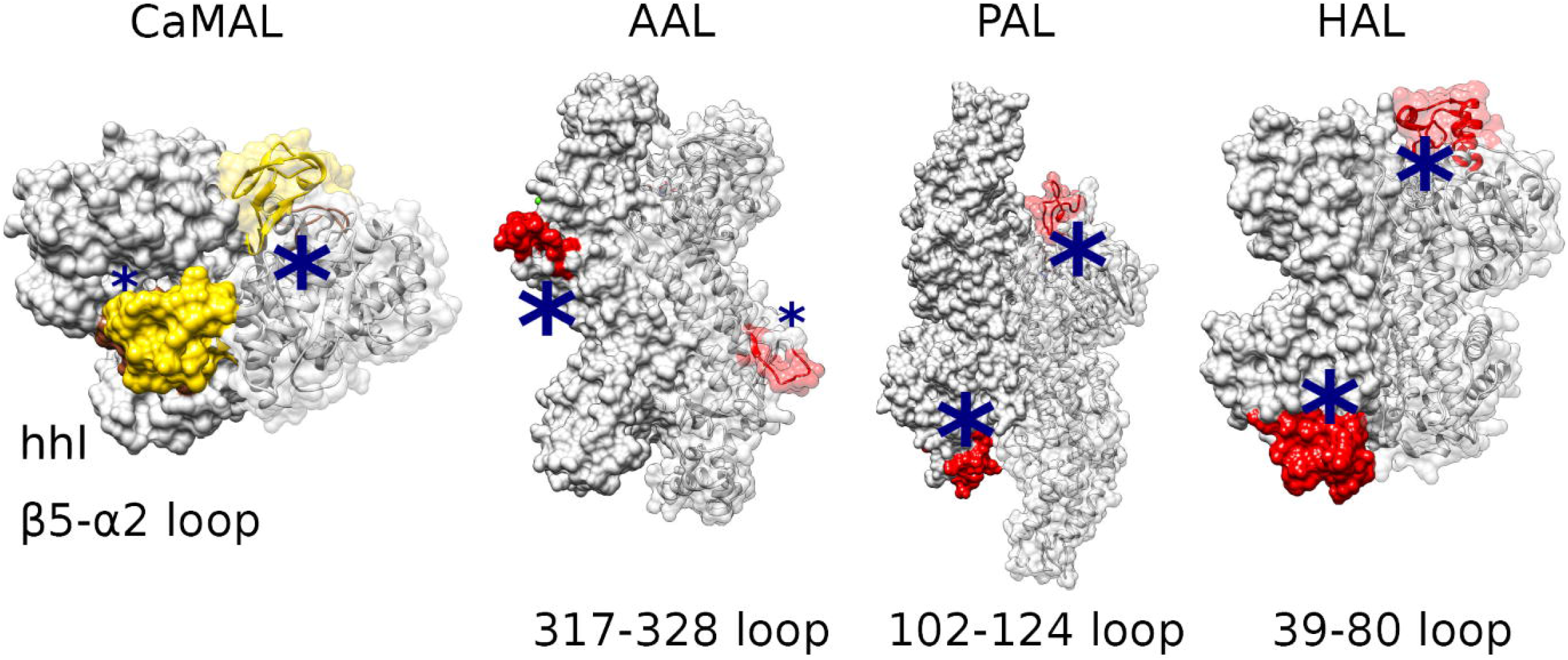
Other ammonia lyases show conformationally heterogeneous structural elements in the proximity of their catalytic sites, which could undergo opening/closing transitions. We showed the structures of: i) CaMAL with the hhl subdomain and the β5-α2 loop highlighted in gold and brown, respectively; ii) aspartate (AAL, PDB entry 3R6V), phenylalanine (PAL, 1W27), and histidine (HAL, 1GKM) ammonia lyases with their disordered regions colored in red in the proximity of the catalytic site, which is indicated by a blue star.

Our results also provide a framework for the future design of mutant variants of CaMAL to entrap the enzyme in its closed state to experimentally validate the effect of the opening/closing transitions on the enzymatic activity of CaMAL. For example, one strategy to pursue could be the engineering of disulphide bridges or mutations that stabilize the interaction networks here identified. A suitable area for mutation is, for example, the region above the catalytic pocket in the proximity of Y240, or the residues G359-S367, which are localized at the side-entrance of the pocket. Our study also opens new directions to understand, at a deeper level, how the substrate recognition and binding happen in CaMAL and how this is regulated by events of conformational changes of the ‘gate’ elements promoted by the substrate.

## Materials and Methods

### CaMAL expression and purification

We retrieved the gene sequence encoding for MAL from *Citrobacter amalonaticus* (CaMAL) (NCBI accession numbers AB005294) from the National Center for Biotechnology Information database and codon optimized for expression in *Escherichia coli* [51]. CaMAL was synthetized by GenScript (Piscataway, NJ, USA) and cloned in the vector pET28b so that a HisTag was introduced in the C-terminal region of the gene. The pET-28b plasmid, which contains the CaMAL gene, was transformed into *Escherichia coli* BL21 (DE3) for protein expression. The cells containing the plasmid were grown in LB media supplemented with 50 μg/ml of kanamycin and autoinduction media with lactose [52] at 30°C at 180 rpm for 16 h. We solubilized the recollected cells in 50 mM Tris-HCl (pH 8) with 300 mM NaCl and 10 mM imidazole and stored at −20°C. The thawed cells were sonicated (Branson Digital Sonifier, model 250) using a maximum amplitude of 30%, in 7 cycles of 30 s and centrifuged for 20 min at 13500 rpms. We purified the supernatants with the expressed protein by immobilized metal ion affinity chromatography (IMAC), using a 1-ml HisTrap column (GE Healthcare, Uppsala, Sweden) and 50 mM phosphate pH 8, 300 mM NaCl, 20mM imidazole buffer. A gradient up to 500 mM imidazole was used to elute the purified protein.

### Size exclusion chromatography experiments

We estimate the molecular weight (MW) of CaMAL by SEC using a Superdex 200 HiLoad 16/600 column (GE Healthcare, Uppsala, Sweden), 50 mM sodium phosphate pH 8.0 with 150 mM sodium chloride as the mobile phase, a flow rate of 1 ml min^−1^, and UV (280 nm) detection. We used blue dextran to determine the column void volume (Vo), which was 43.4 ml. We selected four proteins as molecular mass standards from the gel filtration calibration kit HMW (GE Healthcare, Uppsala, Sweden), i.e. ovalbumin (MW ~ 44 000), conalbumin (MW ~ 75 000), aldolase (MW ~ 158 000) and ferritin (MW ~ 440 000), with elution volumes of 79.8 mL, 73.4 mL, 65.6 mL and 53.4 mL, respectively. The calibration curve was constructed as the partition coefficient (K_av_) *versus* Log (MW). We calculated K_av_ using the equation K_av_ = (V_e_ − V_o_)/ (V_t_ − V_o_), where V_t_ is the total bed volume.

### Preparation of the initial structures for MD simulations

We used the X-ray structure of the homodimer of CaMAL in complex with its natural substrate, i.e. the 3-methyl-aspartic acid (PDB entry 1KKR, [12]) and in the unbound state (PDB entry 1KK0, [12]) as starting structures for the MD simulations. We removed the coordinates of the substrate from the X-ray structure of the bound state of CaMAL. We used Modeller version 9.16 [53] to replace the seleno-methionines in the PDB entries with methionines and we modeled the missing residues at the N- and C-terminal of the protein. In the re-modeling step, we restrained all the protein atoms except for the two residues immediately next to each selenomethionine in the amino acid sequence. We generated 200 different models and we selected the one with the lowest Root Mean Square Deviation (RMSD) calculated on all the methionine atoms using as a reference their original orientation in the X-ray structure. We retained three crystallographic water molecules, which have been suggested to be important for the interaction between the enzyme and its substrate [12].

### MD simulations

We carried out all-atom MD simulations of CaMAL in explicit solvent with *Gromacs* version 5.1.2 [54]. We collected two one-μs MD simulations of CaMAL in its monomeric form, i.e., considering the chain A and chain B separately. Moreover, we run three replicates of the substrate-unbound form CaMAL dimer from the 1KKR structure, using different random seed for the simulations and one replicate starting from the 1KK0 structure to assess the reproducibility of our findings. We employed the CHARMM22 force field with CMAP corrections (namely CHARMM27) [55] and the TIPS3P water model [56]. We predicted the pKa of the histidine residues with PROPKA 3.1 [57] and analyzed the pattern of possible hydrogen bonds for the histidine imidazole nitrogens. We modelled H351 with the hydrogen on the δ nitrogen of the ring, while all the other histidines were modelled with hydrogens on the ε nitrogen.

Dimer simulations were done using a dodecahedral box of water with a minimum distance between protein and box edges of 15 Å applying periodic boundary conditions, while box edges of 30 Å were used for monomers. We used a larger box for the simulations of the monomers to avoid that possible large rearrangements in their structures, which could be caused by the removal of the partner monomer, created artifacts associated with periodic boundary conditions. We gradually equilibrated the system through a series of energy minimization, solvent equilibration, thermalization and pressurization steps using a protocol previously applied to other cases of study [58,59]. In each simulation, we used a 9 Å switch cutoffs for Van der Waals and short-range Coulomb interactions. We employed the Particle-mesh Ewald switch summation method for long-range electrostatic interactions [60]. We stored the conformations every 20 ps, and we used the LINCS algorithm to constrain the heavy atom bond lengths [61]. We collected two one-μs replicates and the third replicate was extended till two μs (see Result and Discussion for more details).

We calculate the minimum distance between the atoms of the protein and its periodic images to identify potential periodic boundary artefacts. In all the monomer simulations the minimal distance was always higher than 33 Å and in the dimer 39 Å.

### Principal Component Analysis

We used a dimensionality reduction technique based on Principal Component Analysis (PCA) [44,45,48] to compare the space sampled by the different MD simulations. The PCA was carried out on a concatenated trajectory of the MD simulations of both monomeric and dimeric states of CaMAL (isolating each monomer from the dimer first) to compare them in the same essential subspace [36,47]. We use the the atoms belonging to the secondary structure of the TIM barrel (i.e., residues 177-187, 209-225, 235-238, 251-265, 270-273, 281-297, 303-306, 313-322, 327-330, 338-351, 355-357, 365-378, and 382-384) for the fitting. We used these residues as a reference for structural superimposition since they are located in the core regions of the TIM barrel domain and do not undergo significant changes in the simulations. We performed the PCA calculations for different subsets of atoms of CaMAL: i) all the protein residues, ii) the region 70-85 (i.e., the β5-α2 loop), and iii) the region 12-51 (i.e., the hhl subdomain) (**Figure 2** and **Figure S2**). We focused on the first two principal components, which account for approximately 43% of the variance in the whole protein scan, and approximately 60% in the β5-α2 loop and hhl subdomain PCA analysis.

### Protein flexibility

We calculated the per-residue Cα-atoms Root Mean Square Fluctuation (RMSF) as an index of flexibility. RMSF was calculated over non-overlapping time windows of 10 ns along the trajectories and then averaged, with a protocol used for other proteins [62–64].

### Changes in secondary structures

We used the DSSP dictionary [65] to estimate the different classes of secondary structures attained for each residue of the β5-α2 loop and hhl subdomain. DSSP defines eight classes of secondary structure elements: 3_10_ helix, α-helix, π-helix, β-sheet, β-bridge, helix turn, bend and coil. For each residue of interest a persistence degree of secondary structure was calculated for each class [66]. The DSSP algorithm is based on calculating the patterns of hydrogen bonds between mainchain carbonyl and amide groups.

### Pairwise distances between selected residues

We calculated the time series of distances between the Cα atom of a reference residue in the binding pocket, i.e. Q329, and the Cα atoms of all the residues of the β5-α2 loop and hhl subdomain.

### Solvent accessible surface area (SASA)

We estimated the SASA for all the atoms of each residues of the catalytic pocket which was either involved in the catalysis or in the interaction with the substrate (i.e., Q73, F170, Q172, H194, Q329, K331, Y356, T360, C361 and L384). We also calculated their cumulative SASA and the total protein SASA per monomer.

### Correlogram

We collected correlogram plots for each trajectory to investigate the coupling between the pairwise selected distances and the different SASA measurements. We used an in-house R script based on the *PerformanceAnalytics, corrplot* and *Hmisc* R packages and the *rcorr()* R function to calculate dependencies between all the pairs of variables. We measured linear dependencies using the Pearson correlation coefficient as a metric for correlation estimate and we calculated the p-value associated to each correlation. We derived the correlograms using the *corrplot()* R function and a hierarchical clustering approach to rank the correlations depending on the degree of couplings between each pair of measurements. We calculated for each measurement (distances and accessibilities) a bivariate scatter plot with the fitted
line and the correlations coefficients with the significance levels at p-values of 0, 0.001, 0.01, 0.05, 0.1, 1. We performed the correlogram analysis for each monomer A and B, separately as derived from the replicates of the CaMAL dimer.

### Correlated motions

We filtered the trajectories on the Cα-atoms and fitted on the Cα-atoms of the same subset of residues used as a reference for the PCA calculation, i.e., belonging to the secondary structure of the TIM barrel (see above). We calculated Cα LMI correlation matrices as previously described [67,68] using the *Wordom* package version 0.22 [69]. We then converted the *Wordom* outputs to the *xPyder* [70] compatible format using the *wordomdccm2dat* tool. LMI values range from 0 (i.e., uncorrelated motions) to 1 (i.e., fully correlated motions). We defined a cutoff of 0.5 to identify significant correlations and remove spurious correlations, as previously applied to other cases of study [58,67]. We also employed a cutoff of distance in the sequence of +/− three residues to filter out the correlations between residues that are next to each other in the primary sequence. We used two non-overlapping time windows of 10 and 100 ns, respectively to estimate average LMI correlation matrices. We calculated the pairwise differences between the LMI matrices calculated for each of the three different dimer replicates, to identify regions in the structure with similar/dissimilar correlations. We compared the LMI matrices calculated for each replicate of the dimer on the monomer A and the monomer B. Moreover, we evaluated the differences between the LMI matrices of the monomer A and the monomer A of different replicate (**Figure S3**). We plotted LMI matrices as heatmaps using an in-house R script based on the ggplot2, reshape2 and viridis R functions.

### Tunnel analyses

We used the CAVER Analyst 1.0 [71] to determine the optimal parameters for the calculations and identification of tunnels along the MD trajectories of CaMAL. We then used the standalone version of CAVER 3.02 software [72] to identify tunnels, which could mediate the access to the active site of CaMAL during the MD simulations. We used as a reference point for the tunnel search the position of the catalytic residues of CaMAL. In particular, we used: i) the nitrogen atom in the amide group of the side chain of Q329; ii) the oxygen atom of the carboxylic group of the side chain of D238; iii) the carbon atom in the methyl group of the side chain of L384 and, iv) the oxygen atom in the hydroxyl group of the side chain of Y356. We performed the tunnel calculations on 1000 representative frames along each trajectory. We removed water molecules and the Mg^2+^ atoms before the calculation. We carried out the tunnel analysis on the MD replicates of the CaMAL dimer, analyzing separately the monomers A and B. We tested different parameters to perform tunnel search and clustering with CAVER, following an optimization procedure previously published [72]. In particular, we selected two set of parameters. The first set aimed to monitor the evolution of solvent-accessible tunnels from the protein surface towards the catalytic site, where we used a probe radius of 1.5 Å, comparable to the size of a water molecule [73], a shell radius of 5 Å, shell depth of 4 Å, and a clustering threshold of 7 Å. In the second set of parameters, we used the same parameters as the first set but with a probe radius of 2.0 Å to resemble the size of the natural substrate 3-methylaspartate. For this purpose, we used as a reference the radius of gyration of 3-methylaspartate (i.e., 2.23 Å). We used for the calculation the conformation of the 3-methylaspartate in the bound crystallographic structure of CaMAL (PDB entry 1KKR [12].

The tunnels were clustered using the hierarchical average-link clustering method, which permit to estimate the dissimilarity (i.e., the distance) of two measured tunnels, dividing each tunnel pathway by sequence of N points starting from the reference site and then evaluating their pairwise distance in their corresponding order. We used PyMOL for visual inspection and graphical representation of the tunnels.

### Contact-based analyses

For the contact-based analyses, we used the CONtact ANalysis (CONAN) software [50]. CONAN permits to obtain a statistical analysis of intramolecular contacts in proteins and to investigate how they evolve along the MD trajectories. We used as cutoffs a r_cut_ value of 10 Å, and r_inter_ and r_high-inter_ values of 5 Å, as previously used [50]. We performed a contact-based cluster analysis using a k-medoid clustering approach based on the RMSD between the contact maps calculated for each frame along each trajectory. The number of maximum clusters was set to five. We observed that CONAN identifies five clusters that are sequentially distributed over the simulation time. We used this analysis to study the temporal evolution of the contacts along the trajectories. We calculated the average distance maps for each cluster, accounting for each pair of residues over all the frames assigned to the same cluster. We analyzed the average distance maps using in-house R scripts. We calculated the percentage increase (or decrease) in the pairwise distances between residues comparing the average contact maps of the first and fifth cluster. We retained as significant only the pairs of residues with a percentage increase in their distance higher than 50%. We analyzed the data using network analysis in *Cytoscape*. The movie of the time-evolution of the contact maps are reported in the Github repository associated to the publication.

### Data and Software Availability

All the software used are freely available. The R and bash scripts, input files for the simulations, input and output files from the modeling step, along with the results generated will be soon freely available in a GitHub repository associated to our publication (https://github.com/ELELAB/MAL_MD). The entire MD trajectories are available upon request.

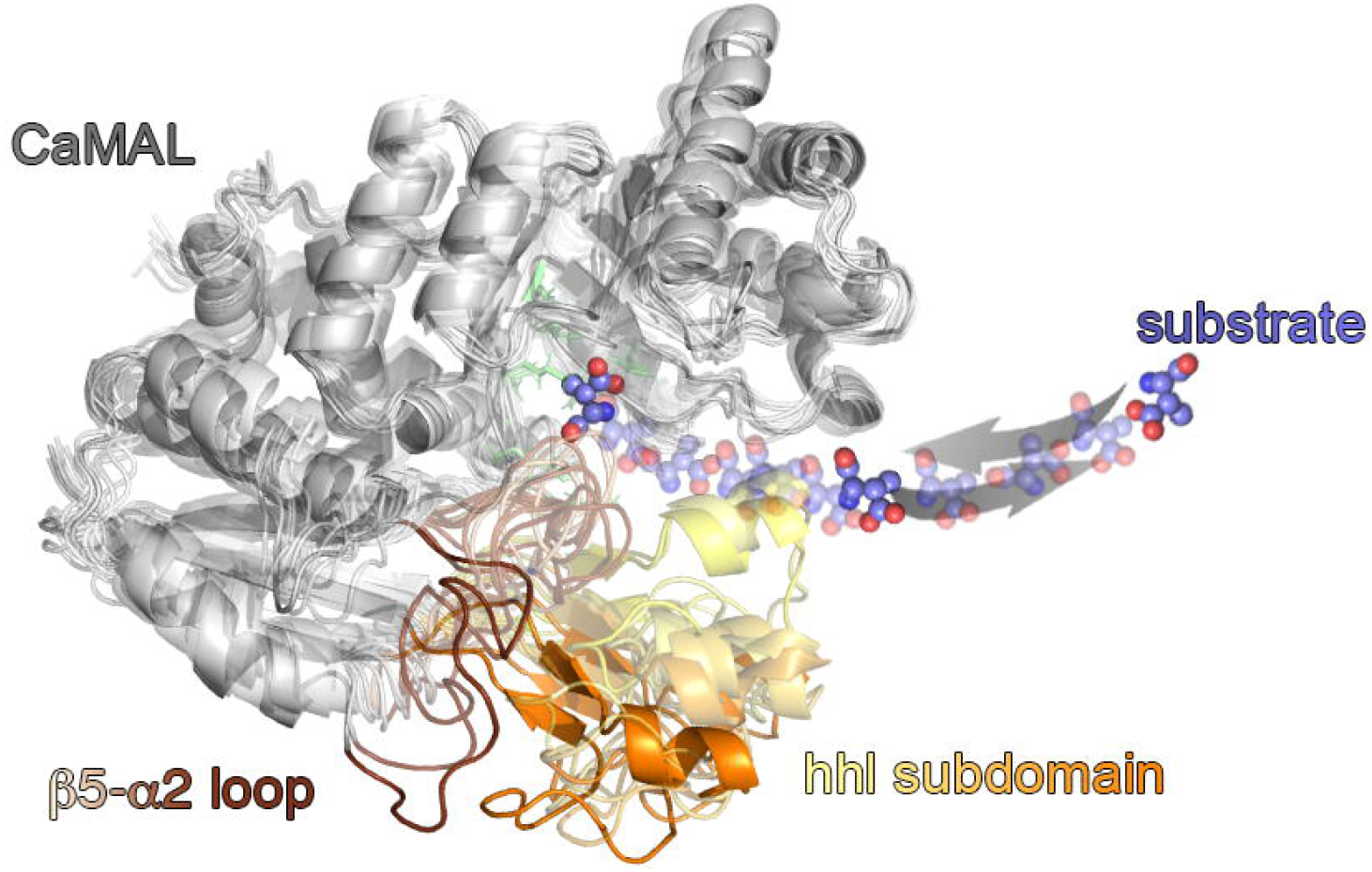

## Acknowledgements

The study was funded by The Swedish Research Council for Environment, Agricultural Sciences and Spatial Planning (FORMAS) (project number 942-2015-1628), the NovoNordisk Foundation under the program for Biotechnology-based Synthesis and Production Research (reference number NNF-17OC0027588) to EP and LO groups and a European Biophysical Society Association (EBSA) bursary granted to VSJ to visit EP group in 2017. EP lab is part of the Center of Excellence in Autophagy, Recycling and Disease (CARD), funded by the Danish National Research Foundation (DNRF125) and also supported by The Danish Council for Independent Research, Natural Science Research Project 1 (102517). The calculations described in this paper were performed using resources provided by the Swedish National Infrastructure for Computing (SNIC) at PDC Center for High Performance Computing under the project SNIC 2017/1-537 and the DeiC National Life Science Supercomputer at DTU.

## Author Contributions

E.P. and M.L. designed the research; E.P. supervised the work; V.S-J. performed the experiments; M.L, Z.S.M. and V.S-J. performed the modeling and simulations; M.L. and Z.S.M. performed the analyses of the simulations; E.P, M.L, Z.S., V.S-J., V.M., L.O. discussed the results; M.L. and Z.S. provided scripts and code for the study; M.L, Z.S. and E.P. wrote the manuscript with suggestions from all the coauthors.

## References

[1] F. Parmeggiani, N.J. Weise, S.T. Ahmed, N.J. Turner, Synthetic and Therapeutic Applications of Ammonia-lyases and Aminomutases, Chem. Rev. (2017) acs.chemrev.6b00824. doi:10.1021/acs.chemrev.6b00824.

[2] M.M. Heberling, B. Wu, S. Bartsch, D.B. Janssen, Priming ammonia lyases and aminomutases for industrial and therapeutic applications, Curr. Opin. Chem. Biol. 17 (2013) 250–260. doi:10.1016/j.cbpa.2013.02.013.

[3] M. De Villiers, V.P. Veetil, H. Raj, J. De Villiers, G.J. Poelarends, Catalytic mechanisms and biocatalytic applications of aspartate and methylaspartate ammonia lyases, ACS Chem. Biol. 7 (2012) 1618–1628. doi:10.1021/cb3002792.

[4] J. Zhang, Y. Liu, A QM/MM study of the catalytic mechanism of aspartate ammonia lyase, J. Mol. Graph. Model. 51 (2014) 113–119. doi:10.1016/j.jmgm.2014.05.003.

[5] A. Heine, G. Herrmann, T. Selmer, F. Terwesten, W. Buckel, K. Reuter, High resolution crystal structure of Clostridium propionicum β-alanyl-CoA: AMmonia lyase, a new member of the “hot dog fold” protein superfamily, Proteins Struct. Funct. Bioinforma. 82 (2014) 2041–2053. doi:10.1002/prot.24557.

[6] K.T. Watts, B.N. Mijts, P.C. Lee, A.J. Manning, C. Schmidt-Dannert, Discovery of a Substrate Selectivity Switch in Tyrosine Ammonia-Lyase, a Member of the Aromatic Amino Acid Lyase Family, Chem. Biol. 13 (2006) 1317–1326. doi:10.1016/j.chembiol.2006.10.008.

[7] N.J. Turner, Ammonia lyases and aminomutases as biocatalysts for the synthesis of α-amino and β-amino acids, Curr. Opin. Chem. Biol. 15 (2011) 234–240. doi:10.1016/j.cbpa.2010.11.009.

[8] A.L. Seff, S. Pilbák, I. Silaghi-Dumitrescu, L. Poppe, Computational investigation of the histidine ammonia-lyase reaction: A modified loop conformation and the role of the zinc(II) ion, J. Mol. Model. 17 (2011) 1551–1563. doi:10.1007/s00894-010-0849-7.

[9] Y. Asano, Y. Kato, C. Levy, P. Baker, D. Rice, Structure and Function of Amino Acid Ammonia-lyases, Biocatal. Biotransformation. 22 (2004) 133–140. doi:10.1080/10242420410001703496.

[10] W.J. Quax, H. Raj, P.G. Tepper, G.J. Poelarends, V. Puthan Veetil, M. de Villiers, F.J. Dekker, Enantioselective Synthesis of N-Substituted Aspartic Acids Using an Engineered Variant of Methylaspartate Ammonia Lyase, ChemCatChem. 5 (2013) 1325–1327. doi:10.1002/cctc.201200906.

[11] H. Raj, W. Szymański, J. De Villiers, H.J. Rozeboom, V.P. Veetil, C.R. Reis, M. De Villiers, F.J. Dekker, S. De Wildeman, W.J. Quax, A.M.W.H. Thunnissen, B.L. Feringa, D.B. Janssen, G.J. Poelarends, Engineering methylaspartate ammonia lyase for the asymmetric synthesis of unnatural amino acids, Nat. Chem. 4 (2012) 478–484. doi:10.1038/nchem.1338.

[12] C.W. Levy, P.A. Buckley, S. Sedelnikova, Y. Kato, Y. Asano, D.W. Rice, P.J. Baker, Insights into enzyme evolution revealed by the structure of methylaspartate ammonia lyase, Structure. 10 (2002) 105–113. doi:10.1016/S0969-2126(01)00696-7.

[13] M. Asuncion, W. Blankenfeldt, J.N. Barlow, D. Gani, J.H. Naismith, The structure of 3-methylaspartase from Clostridium tetanomorphum functions via the common enolase chemical step, J. Biol. Chem. 277 (2002) 8306–8311. doi:10.1074/jbc.M111180200.

[14] H. Raj, B. Weiner, V.P. Veetil, C.R. Reis, W.J. Quax, D.B. Janssen, B.L. Feringa, G.J. Peolarends, Alteration of the diastereoselectivity of 3-methlaspartate ammonia lyase by using structure-based mutagenesis, ChemBioChem. 10 (2009) 2236–2245. doi:10.1002/cbic.200900311.

[15] H. Raj, G.J. Poelarends, The roles of active site residues in the catalytic mechanism of methylaspartate ammonia-lyase, FEBS Open Bio. 3 (2013) 285–290. doi:10.1016/j.fob.2013.07.002.

[16] E. Papaleo, V. Ranzani, F. Tripodi, A. Vitriolo, C. Cirulli, P. Fantucci, L. Alberghina, M. Vanoni, L. De Gioia, P. Coccetti, An Acidic Loop and Cognate Phosphorylation Sites Define a Molecular Switch That Modulates Ubiquitin Charging Activity in Cdc34-Like Enzymes, PLoS Comput. Biol. 7 (2011). doi:10.1371/journal.pcbi.1002056.

[17] E. Papaleo, N. Casiraghi, A. Arrigoni, M. Vanoni, P. Coccetti, L. De Gioia, Loop 7 of E2 Enzymes□: An Ancestral Conserved Functional Motif Involved in the E2-Mediated Steps of the Ubiquitination Cascade, PLoS One. 7 (2012). doi:10.1371/journal.pone.0040786.

[18] S. Pilbák, A. Tomin, J. Rátey, L. Poppe, The essential tyrosine-containing loop conformation and the role of the C-terminal multi-helix region in eukaryotic phenylalanine ammonia-lyases, FEBS J. 273 (2006) 1004–1019. doi:10.1111/j.1742-4658.2006.05127.x.

[19] R.C. Wade, M.E. Davis, B.A. Luty, J.D. Madura, J.A. McCammon, Gating of the active site of triose phosphate isomerase: Brownian dynamics simulations of flexible peptide loops in the enzyme., Biophys. J. 64 (1993) 9–15. doi:10.1016/S0006-3495(93)81335-3.

[20] H.-X. Zhou, S.T. Wlodek, J.A. McCammon, Conformation gating as a mechanism for enzyme specificity, Proc. Natl. Acad. Sci. 95 (2002) 9280–9283. doi:10.1073/pnas.95.16.9280.

[21] M. Kokkinidis, N.M. Glykos, V.E. Fadouloglou, Protein flexibility and enzymatic catalysis, 2012. doi:10.1016/B978-0-12-398312-1.00007-X.

[22] E. Papaleo, G. Saladino, M. Lambrughi, K. Lindorff-Larsen, F.L. Gervasio, R. Nussinov, The role of protein loops and linkers in conformational dynamics and allostery., Chem. Rev. 116 (2016) 6391–6423. doi:10.1021/acs.chemrev.5b00623.

[23] M.K. Rout, B.L. Lee, A. Lin, W. Xiao, L. Spyracopoulos, Active Site Gate Dynamics Modulate the Catalytic Activity of the Ubiquitination Enzyme E2-25K, Sci. Rep. 8 (2018) 1–15. doi:10.1038/s41598-018-25476-8.

[24] A. Gora, J. Brezovsky, J. Damborsky, Gates of Enzymes, Chem. Rev. 113 (2013) 5871–5923. doi:10.1021/cr300384w.

[25] M. Pasi, L. Riccardi, P. Fantucci, L. De Gioia, E. Papaleo, Dynamic properties of a psychrophilic alpha-amylase in comparison with a mesophilic homologue., J. Phys. Chem. B. 113 (2009) 13585–95. doi:10.1021/jp900790n.

[26] D. Tobi, I. Bahar, Structural changes involved in protein binding correlate with intrinsic motions of proteins in the unbound state., Proc. Natl. Acad. Sci. U. S. A. 102 (2005) 18908–13. doi:10.1073/pnas.0507603102.

[27] Z. Kurkcuoglu, A. Bakan, D. Kocaman, I. Bahar, P. Doruker, Coupling between Catalytic Loop Motions and Enzyme Global Dynamics, PLoS Comput. Biol. 8 (2012) e1002705. doi:10.1371/journal.pcbi.1002705.

[28] A. Tousignant, J.N. Pelletier, Protein motions promote catalysis, Chem. Biol. 11 (2004) 1037–1042. doi:10.1016/j.

[29] J.L. Klepeis, K. Lindorff-Larsen, R.O. Dror, D.E. Shaw, Long-timescale molecular dynamics simulations of protein structure and function., Curr. Opin. Struct. Biol. 19 (2009) 120–7. doi:10.1016/j.sbi.2009.03.004.

[30] R.O. Dror, R.M. Dirks, J.P. Grossman, H. Xu, D.E. Shaw, Biomolecular simulation: a computational microscope for molecular biology., Annu. Rev. Biophys. 41 (2012) 429–52. doi:10.1146/annurev-biophys-042910-155245.

[31] T. Saleh, C.G. Kalodimos, Enzymes at work are enzymes in motion, Science (80-.). 355 (2017) 247–248. doi:10.1126/science.aal4632.

[32] E. Papaleo, L. Sutto, F.L. Gervasio, K. Lindorff-Larsen, Conformational changes and free energies in a proline isomerase, J. Chem. Theory Comput. 10 (2014) 4169–4174. doi:10.1021/ct500536r.

[33] R.N. Shinde, S. Karthikeyan, B. Singh, Molecular dynamics studies unravel role of conserved residues responsible for movement of ions into active site of DHBPS, Sci. Rep. 7 (2017) 1–10. doi:10.1038/srep40452.

[34] P. Mereghetti, L. Riccardi, B.O. Brandsdal, P. Fantucci, L. De Gioia, E. Papaleo, Near native-state conformational landscape of psychrophilic and mesophilic enzymes: probing the folding funnel model, J. Phys. Chem. B. 114 (2010) 7609–7619. doi:10.1021/jp911523h.

[35] F. del Caño-Ochoa, A. Grande-García, M. Reverte-López, M. D’Abramo, S. Ramón-Maiques, Characterization of the catalytic flexible loop in the dihydroorotase domain of the human multi-enzymatic protein CAD, J. Biol. Chem. 293 (2018) 18903–18913. doi:10.1074/jbc.RA118.005494.

[36] I. Valimberti, M. Tiberti, M. Lambrughi, B. Sarcevic, E. Papaleo, E2 superfamily of ubiquitin-conjugating enzymes: constitutively active or activated through phosphorylation in the catalytic cleft, Sci. Rep. 5 (2015) 14849. doi:10.1038/srep14849.

[37] E. Papaleo, L. Riccardi, C. Villa, P. Fantucci, L. De Gioia, Flexibility and enzymatic cold-adaptation: A comparative molecular dynamics investigation of the elastase family, Biochim. Biophys. Acta - Proteins Proteomics. 1764 (2006) 1397–1406. doi:10.1016/j.bbapap.2006.06.005.

[38] E. Papaleo, G. Renzetti, Coupled motions during dynamics reveal a tunnel toward the active site regulated by the N-terminal α-helix in an acylaminoacyl peptidase, J. Mol. Graph. Model. 38 (2012) 226–234. doi:10.1016/j.jmgm.2012.06.014.

[39] Y. Kato, Y. Asano, Purification and properties of crystalline 3-methylaspartase from two facultative anaerobes, Citrobacter sp. strain YG-0504 and Morganella morganii strain YG-0601., Biosci. Biotechnol. Biochem. 59 (1995) 93–9. http://www.ncbi.nlm.nih.gov/pubmed/7765982 (accessed March 15, 2019).

[40] E. Papaleo, G. Renzetti, G. Invernizzi, B. Ásgeirsson, Dynamics fingerprint and inherent asymmetric flexibility of a cold-adapted homodimeric enzyme. A case study of the Vibrio alkaline phosphatase, Biochim. Biophys. Acta - Gen. Subj. 1830 (2013) 2970–2980. doi:10.1016/j.bbagen.2012.12.011.

[41] E. Moroni, D.A. Agard, G. Colombo, The Structural Asymmetry of Mitochondrial Hsp90 (Trap1) Determines Fine Tuning of Functional Dynamics, J. Chem. Theory Comput. 14 (2018) 1033–1044. doi:10.1021/acs.jctc.7b00766.

[42] J.M. Flynn, P. Mishra, D.N. a. Bolon, Mechanistic Asymmetry in Hsp90 Dimers, J. Mol. Biol. (2015). doi:10.1016/j.jmb.2015.03.017.

[43] T.H. Kim, P. Mehrabi, Z. Ren, A. Sljoka, C. Ing, A. Bezginov, L. Ye, R. Pomès, R.S. Prosser, E.F. Pai, The role of dimer asymmetry and protomer dynamics in enzyme catalysis, Science (80-.). 355 (2017). doi:10.1126/science.aag2355.

[44] E. Papaleo, P. Mereghetti, P. Fantucci, R. Grandori, L. De Gioia, Free-energy landscape, principal component analysis, and structural clustering to identify representative conformations from molecular dynamics simulations: The myoglobin case, J. Mol. Graph. Model. 27 (2009) 889–899. doi:10.1016/j.jmgm.2009.01.006.

[45] I. Daidone, A. Amadei, Essential dynamics: foundation and applications, Wiley Interdiscip. Rev. Comput. Mol. Sci. 2 (2012) 762–770. doi:10.1002/wcms.1099.

[46] M. Tiberti, E. Papaleo, T. Bengtsen, W. Boomsma, K. Lindorff-Larsen, ENCORE: software for quantitative ensemble comparison, PLoS Comput. Biol. 11 (2015) e1004415.

[47] F. Martín-García, E. Papaleo, P. Gomez-Puertas, W. Boomsma, K. Lindorff-Larsen, Comparing molecular dynamics force fields in the essential subspace., PLoS One. 10 (2015) e0121114. doi:10.1371/journal.pone.0121114.

[48] A. Amadei, A.B. Linssen, H.J. Berendsen, Essential dynamics of proteins., Proteins. 17 (1993) 412–25. doi:10.1002/prot.340170408.

[49] O.F. Lange, H. Grubmüller, Generalized correlation for biomolecular dynamics, Proteins Struct. Funct. Bioinforma. 62 (2005) 1053–1061. doi:10.1002/prot.20784.

[50] D. Mercadante, F. Gräter, C. Daday, CONAN: A Tool to Decode Dynamical Information from Molecular Interaction Maps, Biophys. J. 114 (2018) 1267–1273. doi:10.1016/j.bpj.2018.01.033.

[51] P. Puigbò, E. Guzmán, A. Romeu, S. Garcia-Vallvé, OPTIMIZER: a web server for optimizing the codon usage of DNA sequences., Nucleic Acids Res. 35 (2007) W126–31. doi:10.1093/nar/gkm219.

[52] F.W. Studier, Protein production by auto-induction in high-density shaking cultures, Protein Expr. Purif. 41 (2005) 207–234. doi:10.1016/J.PEP.2005.01.016.

[53] N. Eswar, B. Webb, M. a Marti-Renom, M.S. Madhusudhan, D. Eramian, M.-Y. Shen, U. Pieper, A. Sali, Comparative protein structure modeling using MODELLER., Curr. Protoc. Protein Sci. Chapter 2 (2007) Unit 2.9. doi:10.1002/0471140864.ps0209s50.

[54] M.J. Abraham, T. Murtola, R. Schulz, S. Páll, J.C. Smith, B. Hess, E. Lindahl, GROMACS: High performance molecular simulations through multi-level parallelism from laptops to supercomputers, SoftwareX. 2 (2015) 19–25. doi:10.1016/j.softx.2015.06.001.

[55] P. Bjelkmar, P. Larsson, M.A. Cuendet, B. Hess, E. Lindahl, Implementation of the CHARMM force field in GROMACS: analysis of protein stability effects from correction Maps, virtual interaction sites, and water models., J. Chem. Theory Comput. 6 (2010) 459–66. doi:10.1021/ct900549r.

[56] W.L. Jorgensen, J. Chandrasekhar, J.D. Madura, R.W. Impey, M.L. Klein, Comparison of simple potential functions for simulating liquid water, J. Chem. Phys. 79 (1983) 926. doi:10.1063/1.445869.

[57] M.H.M. Olsson, C.R. Søndergaard, M. Rostkowski, J.H. Jensen, PROPKA3: Consistent Treatment of Internal and Surface Residues in Empirical p K a Predictions, J. Chem. Theory Comput. 7 (2011) 525–537. doi:10.1021/ct100578z.

[58] M. Tiberti, G. Invernizzi, E. Papaleo, (Dis)similarity Index To Compare Correlated Motions in Molecular Simulations, J. Chem. Theory Comput. 11 (2015) 4404–4414. doi:10.1021/acs.jctc.5b00512.

[59] M. Nygaard, T. Terkelsen, A.V. Olsen, V. Sora, J. Salamanca, F. Rizza, S. Bergstrand, M. Di Marco, M. Vistesen, M. Lambrughi, M. Jaattela, T. Kallunki, E. Papaleo, The mutational landscape of the oncogenic MZF1 SCAN domain in cancer, Front. Mol. Biosci. 3 (2016) 1–18. doi:10.3389/fmolb.2016.00078.

[60] U. Essmann, L. Perera, M.L. Berkowitz, T. Darden, H. Lee, L.G. Pedersen, A smooth particle mesh Ewald method, J. Chem. Phys. 103 (1995) 8577. doi:10.1063/1.470117.

[61] B. Hess, H. Bekker, H. Berendsen, J. Fraaije, LINCS: A linear constraint solver for molecular simulations, J. Comput. Chem. 12 (1993) 1463–1472. doi:10.1002/(SICI)1096-987X(199709)18:12<1463::AID-JCC4>3.0.CO;2-H.

[62] K.R. Óskarsson, M. Nygaard, B. Ellertsson, S.H. Thorbjarnardottir, E. Papaleo, M.M. Kristjánsson, A single mutation Gln142Lys doubles the catalytic activity of VPR, a cold adapted subtilisin-like serine proteinase, Biochim. Biophys. Acta - Proteins Proteomics. 1864 (2016) 1436–1443. doi:10.1016/j.bbapap.2016.07.003.

[63] A. Di Rita, A. Peschiaroli, P. D’Acunzo, D. Strobbe, Z. Hu, J. Gruber, M. Nygaard, M. Lambrughi, G. Melino, E. Papaleo, J. Dengjel, S. El Alaoui, M. Campanella, V. Dötsch, V. V. Rogov, F. Strappazzon, F. Cecconi, HUWE1 E3 ligase promotes PINK1/PARKIN-independent mitophagy by regulating AMBRA1 activation via IKKα, Nat. Commun. 9 (2018) 3755. doi:10.1038/s41467-018-05722-3.

[64] D. Michetti, B.O. Brandsdal, D. Bon, G.V. Isaksen, M. Tiberti, E. Papaleo, A comparative study of cold- and warm-adapted Endonucleases A using sequence analyses and molecular dynamics simulations, PLoS One. 12 (2017) e0169586. doi:10.1371/journal.pone.0169586.

[65] W. Kabsch, C. Sander, Dictionary of protein secondary structure: Pattern recognition of hydrogen-bonded and geometrical features, Biopolymers. 22 (1983) 2577–2637. doi:10.1002/bip.360221211.

[66] E. Papaleo, M. Pasi, M. Tiberti, L. De Gioia, Molecular dynamics of mesophilic-like mutants of a cold-adapted enzyme: insights into distal effects induced by the mutations., PLoS One. 6 (2011) e24214. doi:10.1371/journal.pone.0024214.

[67] G. Invernizzi, M. Tiberti, M. Lambrughi, K. Lindorff-Larsen, E. Papaleo, Communication Routes in ARID Domains between Distal Residues in Helix 5 and the DNA-Binding Loops, PLoS Comput. Biol. 10 (2014) e1003744. doi:10.1371/journal.pcbi.1003744.

[68] O.F. Lange, H. Grubmüller, Full correlation analysis of conformational protein dynamics., Proteins. 70 (2008) 1294–312. doi:10.1002/prot.21618.

[69] M. Seeber, A. Felline, F. Raimondi, S. Muff, R. Friedman, F. Rao, A. Caflisch, F. Fanelli, Wordom: a user-friendly program for the analysis of molecular structures, trajectories, and free energy surfaces., J. Comput. Chem. 32 (2011) 1183–94. doi:10.1002/jcc.21688.

[70] M. Pasi, M. Tiberti, A. Arrigoni, E. Papaleo, xPyder: a PyMOL plugin to analyze coupled residues and their networks in protein structures., J. Chem. Inf. Model. 279 (2012) 1–6. doi:10.1021/ci300213c.

[71] B. Kozlikova, E. Sebestova, V. Sustr, J. Brezovsky, O. Strnad, L. Daniel, D. Bednar, A. Pavelka, M. Manak, M. Bezdeka, P. Benes, M. Kotry, A. Gora, J. Damborsky, J. Sochor, CAVER Analyst 1.0: graphic tool for interactive visualization and analysis of tunnels and channels in protein structures, Bioinformatics. 30 (2014) 2684–2685. doi:10.1093/bioinformatics/btu364.

[72] E. Chovancova, A. Pavelka, P. Benes, O. Strnad, J. Brezovsky, B. Kozlikova, A. Gora, V. Sustr, M. Klvana, P. Medek, L. Biedermannova, J. Sochor, J. Damborsky, CAVER 3.0: A Tool for the Analysis of Transport Pathways in Dynamic Protein Structures, PLoS Comput. Biol. 8 (2012) e1002708. doi:10.1371/journal.pcbi.1002708.

[73] M. Manak, M. Zemek, J. Szkandera, I. Kolingerova, E. Papaleo, M. Lambrughi, Hybrid Voronoi diagrams, their computation and reduction for applications in computational biochemistry, J. Mol. Graph. Model. 74 (2017) 225–233. doi:10.1016/j.jmgm.2017.03.018.

