## Supplementary material for "Conformational gating in ammonia lyases": Figure S1

**Figure S1**. Per-residue RMSF profile of monomer A and B MD simulations in comparison to the corresponding monomer in the replicate 1 of the CaMAL MD simulation of the dimeric form.

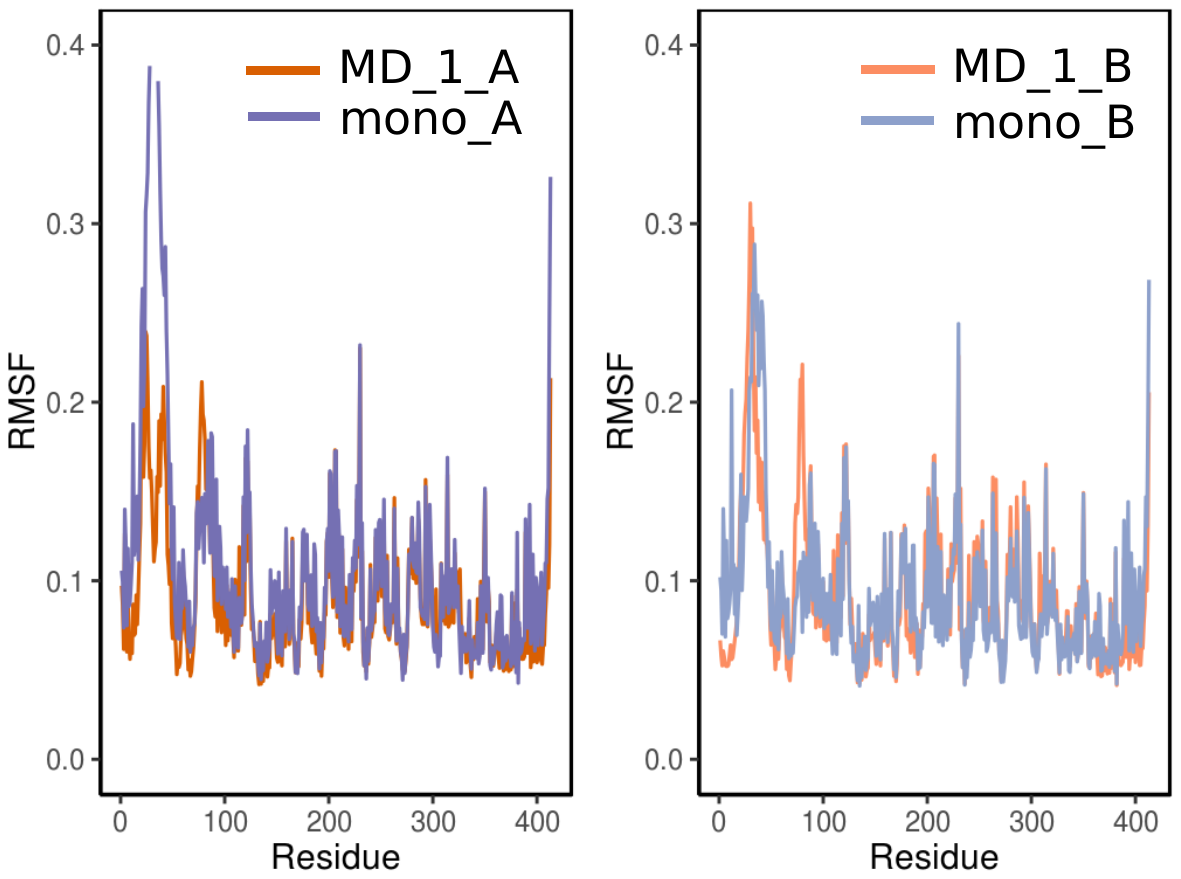
