## Supplementary material for "Conformational gating in ammonia lyases": Figure S2

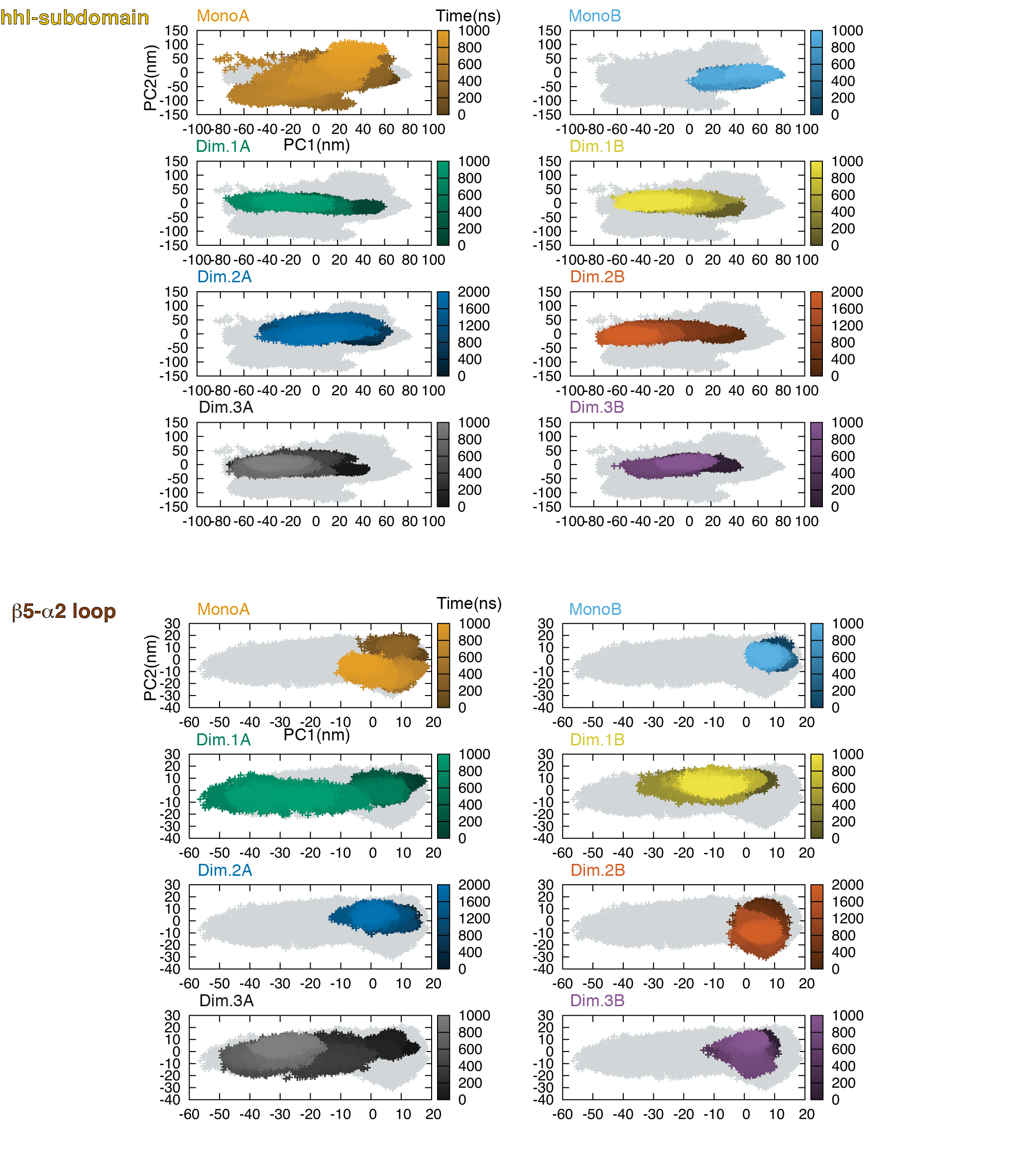
**Figure S2. Flexibility of the hhl subdomain (upper panel) and the β5-α2 loop (lower panel) from Principal Component Analysis (PCA) analysis performed considering only the atoms comprised in each of these two regions of CaMAL.** The ensemble sampled by each trajectory on the 2D distribution plot from the PCA analysis is shown, pointing out the different conformational states collected by the two trajectories of the separated monomers of CaMAL (monomer A in orange and monomer B in light blue) compared to the three replicates of the dimeric form of CaMAL (replicate 1 monomer A in green and monomer B in yellow, replicate 2 monomer A in dark blue monomer B in red, replicate 3 monomer A in grey and monomer B in purple). The time evolution of each trajectory in the conformational space is represented as shade of color, from dark (0 ns) to light (end of the trajectory).
