## Supplementary material for "Conformational gating in ammonia lyases": Figure S3

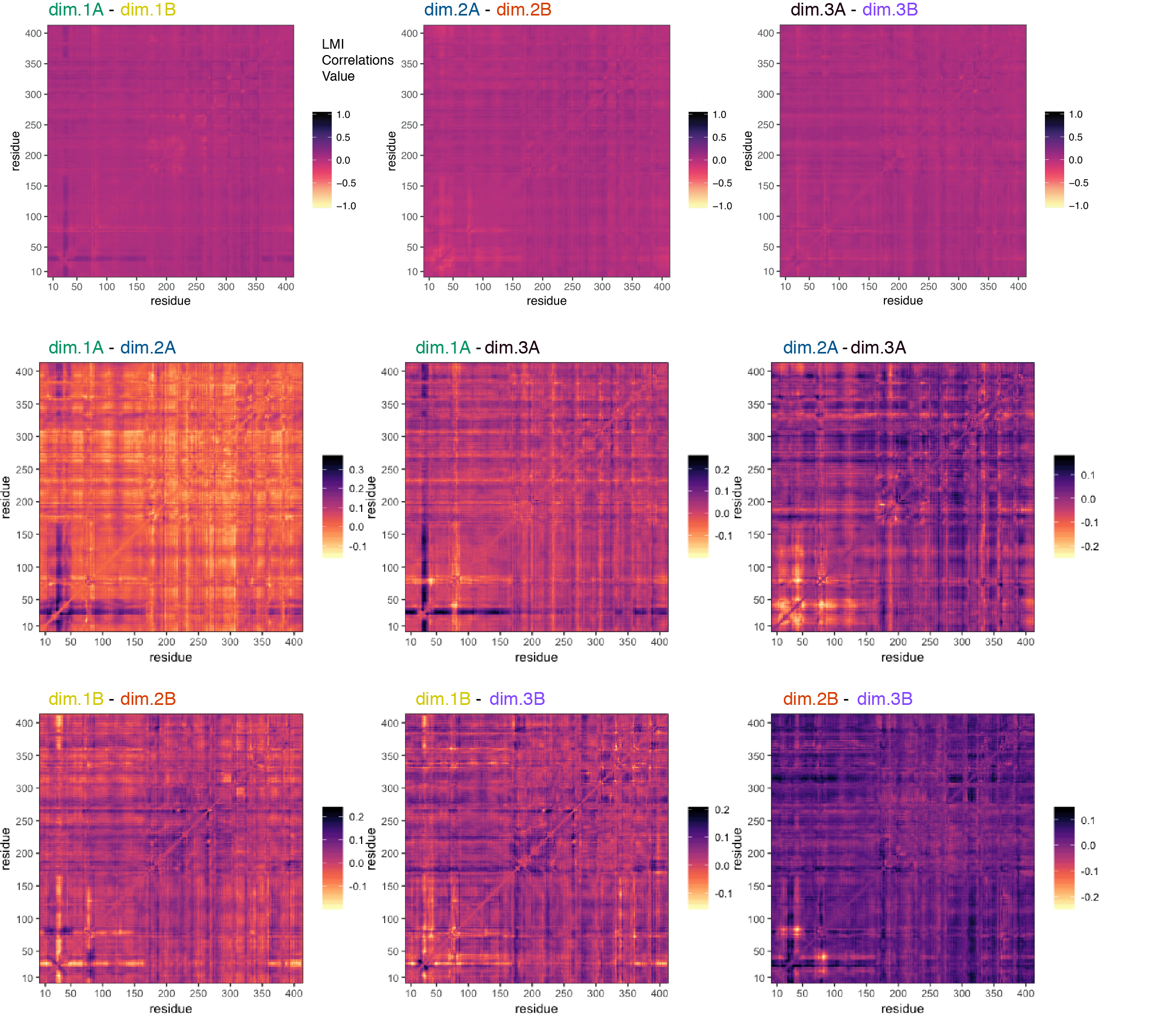

**Figure S3. Comparison of the LMI matrices calculated on the three CaMAL dimer replicates and averaged over 100ns time-windows.** We calculated the pairwise differences between the LMI matrices calculated for the monomer A and the monomer B of the same replicate (upper first three panels for replicate 1, 2 and 3 respectively) as well as the differences between the LMI matrices of the monomer A and the monomer A (middle three panels) and of the monomer B and the monomer B (bottom three panels) of different replicates. The LMI matrices calculated are consistent along all of replicate, describing a similar pattern of correlated fluctuations between the residues for both the monomer A and monomer B, with differences in the range of +/-0.3.
